# Diverse but unique astrocytic phenotypes during embryonic stem cell differentiation, culturing and aging

**DOI:** 10.1101/2021.08.02.454573

**Authors:** Kiara Freitag, Pascale Eede, Andranik Ivanov, Shirin Schneeberger, Tatiana Borodina, Sascha Sauer, Dieter Beule, Frank L. Heppner

## Abstract

Astrocytes are resident glia cells of the central nervous system (CNS) that play complex and heterogeneous roles in brain development, homeostasis and disease. Since their vast involvement in health and disease is becoming increasingly recognized, suitable and reliable tools for studying these cells *in vivo* and *in vitro* are of utmost importance. One of the key challenges hereby is to adequately mimic their context-dependent *in vivo* phenotypes and functions *in vitro*. To better understand the spectrum of astrocytic variations in defined settings we performed a side-by-side-comparison of embryonic stem cell (ESC)-derived astrocytes as well as primary neonatal and adult astrocytes, revealing major differences on a functional and transcriptomic level, specifically on proliferation, migration, calcium signaling and cilium activity. Our results highlight the need to carefully consider the choice of astrocyte origin and phenotype with respect to age, isolation and culture protocols based on the respective biological question.

## Introduction

Astrocytes are central nervous system (CNS)-resident glia cells that have long been considered as passive supporting cells, yet they have recently caught major attention due to their involvement in many neurological diseases. Astrocytes form vast intracellular and intercellular networks within the brain that influence homeostatic cell metabolism and function ^1,2^. As the main constituent of the neurovascular junction, astrocytes ensure the energy supply to the brain ^3,4,5^ whilst also controlling neuronal health by recycling neurotransmitters and by promoting the formation of neural networks ^6^. Conversely, due to their crucial role in brain homeostasis, dysfunction of astrocytes has a substantial impact on brain disorders, in particular upon neurodegenerative and neuroinflammatory diseases ^7^.

With an increasing need to properly assess astrocytes, the need for efficient isolation and culturing protocols keeping astrocytes close to their *in vivo* profile rises. Many methods to isolate and study murine astrocytes have been established, yet such approaches need to be carefully selected based on the underlying scientific question ^8^. *In vitro* approaches using primary astrocytes provide a useful platform to dissect specific astrocytic functions and molecular mechanisms, however maintaining an *in vivo*-like phenotype has been a major challenge. It is widely known that the classic astrocyte isolation technique using postnatal rodent brains based on McCarthy and de Vellis ^9^ induces a reactive astrocytic phenotype ^10^, thus changing the presumed *in vivo* profile of astrocytes. Similarly, astrocytic developmental stages along with their functional and transcriptomic differences are often not taken into account or compared between isolation techniques. Consequently, many studies assessing astrocyte functions only rely on cultures from neonatal astrocytes, thus neglecting the major functional changes that occur context-dependently during development and aging ^11,12,13^.

To generate a baseline reference of transcriptomic and functional changes of various astrocyte culture settings and origins, we compared magnetic activated cell sorted (MACS) ACSA-2-positive murine astrocytes ^14,15^ isolated from (1) neonatal and from (2) adult wild-type mice to (3) astrocytes generated from embryonic stem cells (ESC) (AGES) ^16^. So far, thorough side-by-side comparisons of ESC- or induced pluripotent stem cell (iPSC)-derived astrocytes to their primary adult and neonatal counterparts taking phenotypic, functional and transcriptomic characteristics into account are to our knowledge lacking. Additionally, we investigated how cellular properties of neonatal and adult astrocytes change upon culturing in order to highlight which *in vivo* functions are specifically altered in an *in vitro* setting. Analysis of astrocyte markers, transcriptomic profiles and functional properties revealed major differences between the various astrocyte populations. Whilst functions related to trophic support such as synaptic vesicle transport and dendritic spine development were lost upon culturing of primary astrocytes, key age-specific differences in cilium expression were retained. AGES displayed distinctive transcriptomic and functional signatures resembling astrocytic characteristics that did not fully align with primary astrocyte profiles and most likely represent an intermediate state between primary cells and neural stem cells (NSCs). These data highlight the importance of carefully aligning experimental requirements to the underlying biological question when assessing astrocytic properties *in vitro* and *ex vivo*.

## Results & Discussion

### Distinct astrocytic marker profiles between AGES, cultured and directly isolated primary astrocytes

Given their vital functions in health and disease, it is essential to have valid and robust astrocyte *in vitro* models allowing to assess their particular contributions in physiological as well as CNS disease settings. Therefore, we compared the protein and transcriptomic profiles of three widely used astrocytic cell types, namely of cultured and freshly isolated neonatal and adult astrocytes as well as AGES. Whole hemisphere neonatal astrocytes were isolated from four- to eight-day-old C57BL/6J wild type mice by MACS using the neural tissue dissociation kit (NTDK) and anti-ACSA-2 magnetic microbeads (Fig. 1A). For adult astrocyte populations, cells were isolated from > 100-day old C57BL/6J mice also using MACS, with the only change that the adult brain dissociation kit (ABDK) including debris and red blood cell removal steps was used (Fig. 1B). Cultured cells were used for downstream analyses after 7-10 days *in vitro*. For evaluating how efficiently AGES may replace primary astrocytes for the purpose of *in vitro* studies, murine ESCs (mESC) were differentiated into NSCs. After reaching purity (Fig. S2A), NSCs were terminally differentiated into AGES within three to five days by adding bone morphogenetic protein 4 (BMP4) ^16^ (Fig. 1C). Fluorescence activated cell sorting (FACS) for the astrocyte-specific cell surface marker ACSA-2 indicated that directly isolated astrocytes showed the highest level of purity (97-98.7 %), which was slightly reduced upon culturing and in AGES (Fig. S1A-F). Due to high purity levels in all conditions, we consider MACS as a suitable method for the isolation of pure neonatal and adult astrocyte populations. As ACSA-2 is also expressed by a few non-astrocytic cell types such as glial progenitor cells, neural stem cells and radial glia ^15, 17^, we extended our purity analysis by assessing a wide range of established astrocytic markers. Staining for the astrocytic intermediate filament proteins GFAP and Nestin as well as the astrocytic glutamate transporters GLAST and GLT-1 revealed that all astrocytic cells expressed these markers (Fig. 1D-F). Gene expression analysis of astrocytic cell markers showed increased *Gfap* expression levels in cultured primary cells and in AGES (Fig. 1G). However, GFAP expression can vary greatly between astrocytic cell populations, physical activity and reactive states ^18^, highlighting the need to include a wider range of astrocytic cell markers ^19^. Compared to directly snap-frozen neonatal astrocytes, expression of *Aldh1l1* was reduced in cultured cells and hardly seen in AGES (Fig. 1H). In contrast, *Slc1a3* (Glast) levels were highest in directly isolated neonatal and adult astrocytes and similar in all three cultured cell types (Fig. 1I). Neonatal and adult astrocyte cultures therefore mimic previously described maturation-dependent gene expression patterns ^13,20^, whilst AGES do not fully reflect the marker profile of primary neonatal or adult astrocytes which could also be influenced by the differing culturing conditions. Gene expression analysis of non-astrocytic cell markers revealed that presence of microglia and oligodendrocytes can vary with the age and culturing of astrocytes (Fig. S1G-J). When characterizing astrocyte cultures, a wide range of astrocytic and non-astrocytic markers should therefore be used.

**Figure 1:**
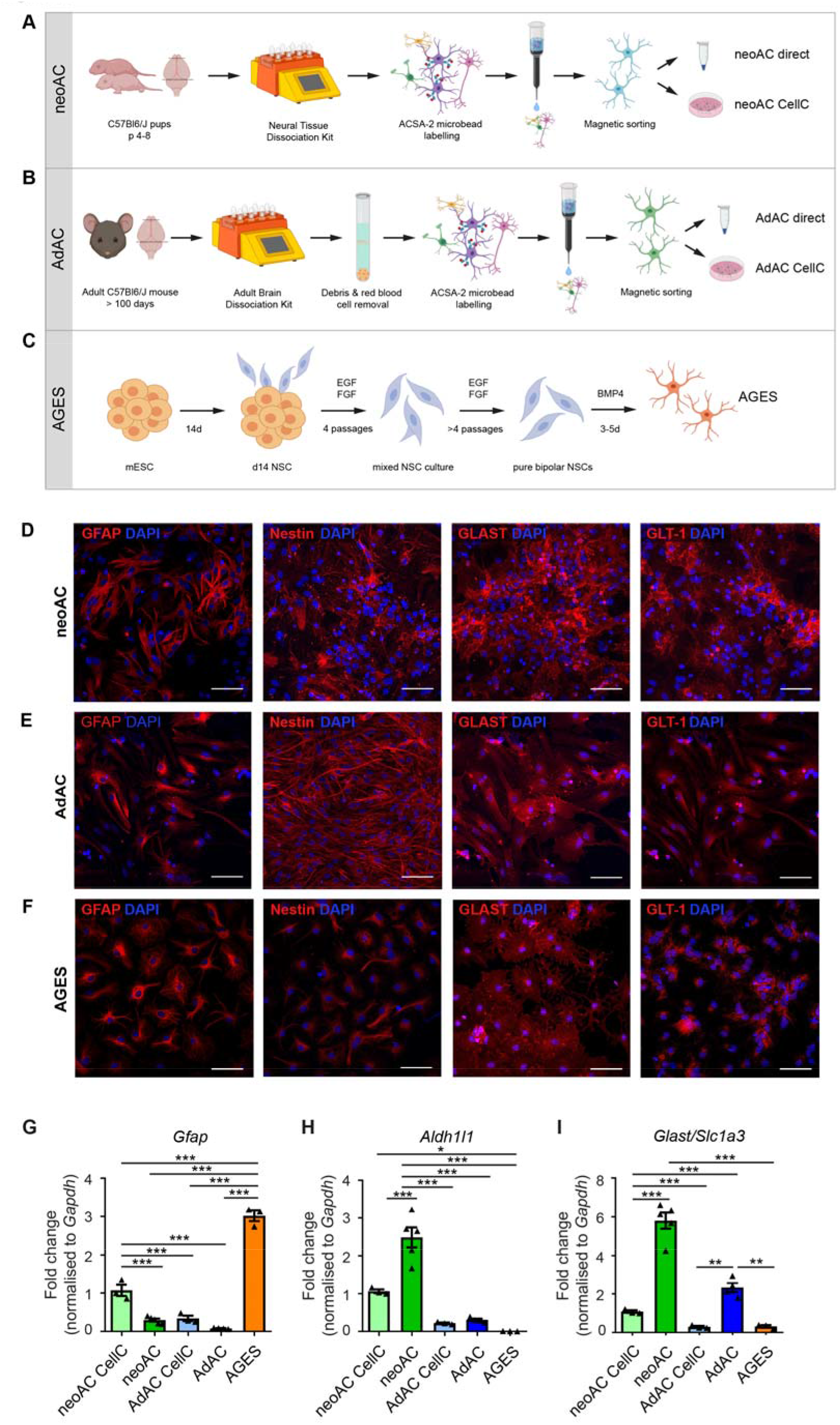
Distinct astrocytic marker profiles between AGES, cultured and directly isolated primary astrocytes. (**A**) Neonatal astrocytes (neoAC) were isolated from four- to eight-day-old C57BL/6J mice by magnetic activated cell sorting (MACS) using the neural tissue dissociation kit (NTDK) and anti-ACSA-2 magnetic microbeads. Cells were either snap-frozen (neoAC direct) or cultured (neoAC CellC). (**B**) Adult astrocytes (AdAC) were isolated from >100d C57BL/6J mice by MACS using the adult brain dissociation kit (ABDK) including a debris removal and red blood cell removal step followed by ACSA-2 microbead labelling. Cells were either snap-frozen (AdAC direct) or cultured (AdAC CellC). (**C**) Astrocytes generated from embryonic stem cells (AGES) were differentiated from mouse embryonic stem cells (mESCs) by creating neural stem cells (NSCs). For receiving pure bipolar NSC cultures, NSCs were cultured for at least eight passages in medium containing the growth factors EGF and FGF. Differentiation into AGES was induced by adding bone morphogenetic protein 4 (BMP4) for three to five days. (**D-F**) Fluorescent immunocytochemistry of (**D**) cultured neonatal astrocytes, (**E**) cultured adult astrocytes and (**F**) AGES was performed for the astrocytic marker proteins: glial fibrillary acidic protein (GFAP), Nestin and the astrocytic glutamate transporters GLAST and GLT-1. Images were taken by confocal microscopy. Scale bar = 50 µm. (**G-I**) Gene expression of the astrocytic markers (**G**) *Gfap* (*\*\*\*P* < 0.001), **(H)** *Aldh1l1* (*\*P* = 0.0130; *\*\*\*P* < 0.001) and (**I**) *Slc1a3*/Glast (AdAC CellC vs AdAC direct *\*\*P* = 0.0034; AdAC direct vs AGES *\*\*P* = 0.0037; *\*\*\*P* < 0.001) was determined by quantitative real-time PCR. All expression values were normalized to the internal control *Gapdh* and cultured neonatal astrocytes as a reference. Mean ± SEM; neoAC CellC (n=3), neoAC (n=5), AdAC CellC (n=3), AdAC (n=4), AGES (n=3); ANOVA with Tukey’s post hoc test.

### Similarities in glucose uptake, lactate release and synaptosome uptake oppose differences in proliferative, migratory and calcium signaling in AGES and astrocyte cultures

To investigate whether the differences seen in the astrocytic marker profile also implicate differences in functional properties, we assessed the maintenance of homeostatic functions of astrocytes in culture. Proliferation in astrocytes is known to cease with ageing ^21^, which we confirmed in an 5-ethynyl-2’-deoxyuridine (EdU)-based assay, where cultured adult astrocytes proliferated less than neonatal astrocytes, while AGES, as terminally-differentiated cells, showed a proliferation rate similar to adult astrocytes (Fig. 2A) which was not influenced by the amount of dying cells (Fig. 2B). To mimic wound healing upon tissue injury, a major feature of astrocytes *in vivo*, a confluent astrocyte layer was disrupted by creating a wound gap *in vitro*, showing that adult astrocytes migrate or sense wound gaps significantly faster than neonatal astrocytes and AGES, with AGES exerting a very slow response rate (Fig. 2C-F, Fig. S2B-C). Assessing further physiological functions of astrocytes such as metabolism of glucose and lactate ^22^ as well as synapse elimination ^23^ revealed no differences between primary cultured astrocytes and AGES (Fig. 2G-I), emphasizing that AGES are a suitable *in vitro* model for assessing these physiological functions of astrocytes. To investigate calcium signaling as the main communication system of astrocytes ^24^ we performed live imaging of astrocytes incubated with the calcium indicator Fluo-4, which is coupled with acetoxymethyl (AM) to ensure fluorescence only after cell entry (Fig. 2J-N). We found that AGES responded fastest to ATP stimulation (Fig. 2L) and their maximum fluorescent intensity i.e. the amount of calcium released, was highest compared to neonatal and adult astrocytes (Fig. 2M), which is in line with previous reports showing that the amplitude of the spontaneous calcium spike was significantly higher in human iPSC-astrocytes compared to primary astrocytes ^25^. No differences in the duration of the calcium response were seen between the cell types (Fig. 2N). Taken together, key differences in functions of AGES compared to primary astrocyte cultures were found in proliferation, response to wound gaps and calcium release properties. Notably, it was suggested that even in instances of damage, cell migration might rather be a non-physiological function of astrocytes *in vivo* and may represent a behavior only acquired in culture system *in vitro* ^26^. Despite showing differences to primary astrocytes, AGES might therefore be more suitable in modelling the *in vivo-*like migration behavior of astrocytes. Thus, it is of utmost relevance to consider the identified functional differences when choosing a model system for a respective biological investigation.

**Figure 2:**
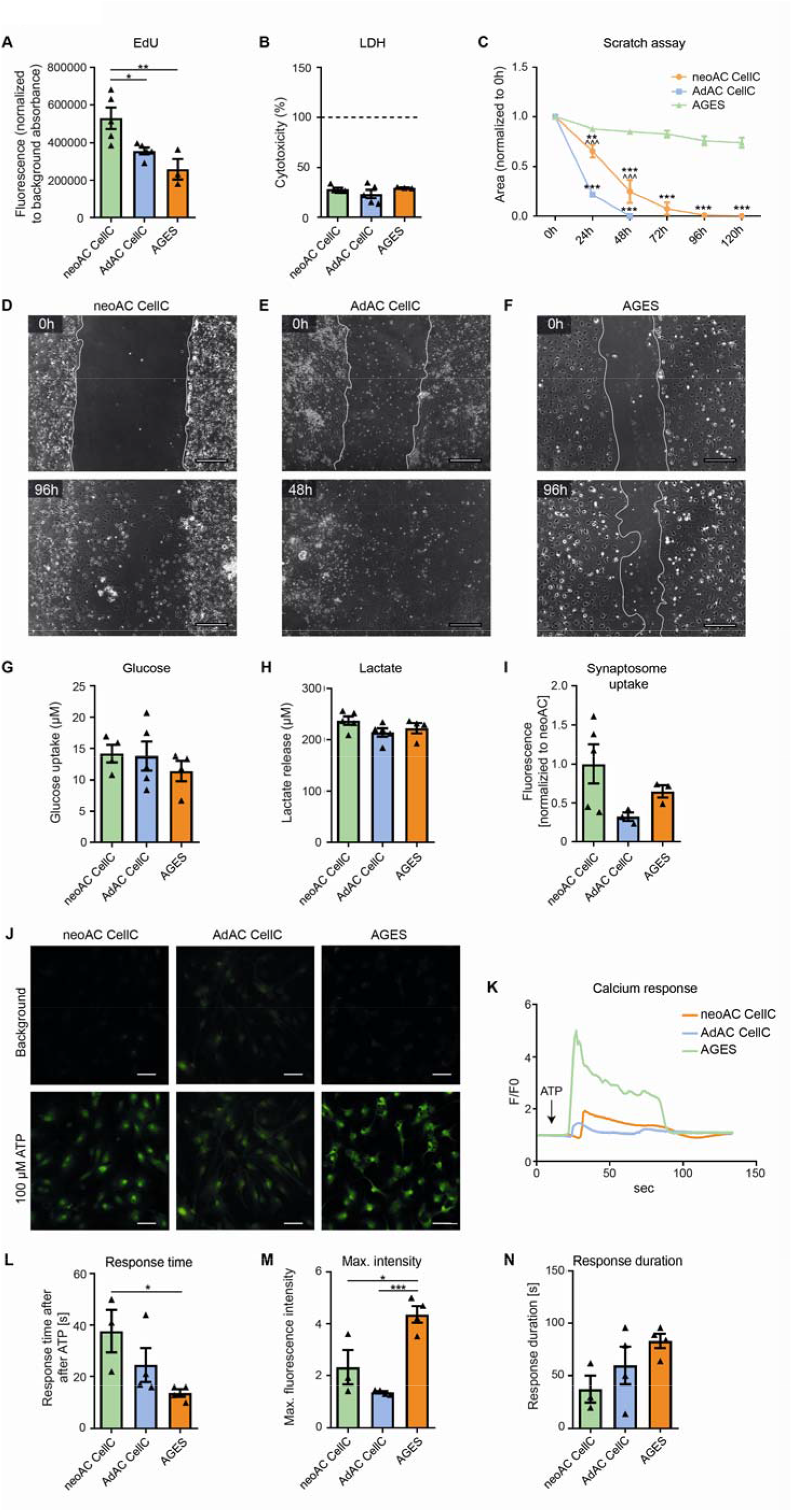
Similarities in glucose uptake, lactate release and synaptosome uptake oppose differences in proliferative, migratory and calcium signaling of AGES and isolated astrocytes. (**A**) Proliferation was measured by fluorescent immunolabeling for the thymidine analogue EdU which is incorporated during cell division. Fluorescent signal was normalized to background absorbance. ANOVA with Tukey’s post hoc test *\*P* = 0.0365; *\*\*P* = 0.0068 ***(B***) Cytotoxicity was measured by determining the levels of lactate dehydrogenase (LDH) release to the cell supernatant. As a positive control, each cell type was lysed with Triton-X to induce maximal LDH release per cell (= 100 %, dashed line). Medium served as a negative control and all values were plotted as percentages of the maximal LDH release. Mean ± SEM; neoAC CellC (n=5), AdAC CellC (n=5), AGES (n=3). ANOVA with Tukey’s post hoc test *P >* 0.05. (**C**) A scratch assay was used to determine the migratory behaviour of cultured neonatal astrocytes (neoAC CellC), cultured adult astrocytes (AdAC CellC) and AGES. The area of the wound gap normalized to time point 0 h is shown over time. * significance compared to AGES; ^ significance compared to AdAC CellC. Mean ± SEM; neoAC CellC (n=4), AdAC CellC (n=4), AGES (n=4); Two-way ANOVA with Bonferroni post hoc test; *\*\*P* = 0.004; *\*\*\*P* < 0.001; *^^^P* < 0.001. (**D-F**) Representative phase contrast images of the wound gap at the starting point (0 h) and after 48 h/96 h are shown of (**D**) cultured neonatal astrocytes (neoAC CellC), (**E**) cultured adult astrocytes (AdAC CellC) and (**F**) AGES. Scale bar = 200 µm. (**G**) Glucose uptake (in µM) in cultured neonatal astrocytes (neoAC CellC), cultured adult astrocytes (AdAC CellC) and AGES was measured with a luminescence-based assay. Mean ± SEM; neoAC CellC (n=4), AdAC CellC (n=5), AGES (n=4). ANOVA with Tukey’s post hoc test *P >* 0.05. (**H**) To determine the lactate released by each cell type (in µM), lactate was measured in the cell supernatant with a luminescence-based assay. Mean ± SEM; neoAC CellC (n=5), AdAC CellC (n=5), AGES (n=4). ANOVA with Tukey’s post hoc test *P >* 0.05. (**I**) pH-sensitive fluorescently labelled synaptosomes isolated from C57Bl/6J mice were cocultured with cultured neonatal astrocytes (neoAC CellC), cultured adult astrocytes (AdAC CellC) and AGES. The fluorescence was measured and represents the uptake of synaptosomes. The fluorescent signals were normalized to cultured neonatal astrocytes. Mean ± SEM; neoAC CellC (n=5), AdAC CellC (n=3), AGES (n=3). ANOVA with Tukey’s post hoc test *P >* 0.05. (**J-N**) Calcium signaling of all three cell types was determined by calcium imaging using Fluo-4, AM as a calcium indicator and 100 µM ATP as a stimulus for calcium release. (**J**) Representative images of cultured neonatal astrocytes (neoAC CellC), cultured adult astrocytes (AdAC CellC) and AGES are shown before ATP stimulation (= background) and after ATP stimulation. Scale bar = 50 µm. (**K**) The fluorescence intensity normalized to background fluorescence (F/F0) is shown over time for each cell type. (**L**) The time until cells responded with a calcium peak was measured. ANOVA with Tukey’s post hoc test *\*P* = 0.0493 (**M**) The maximum fluorescence intensity was compared between all cell types. ANOVA with Tukey’s post hoc test *\*P* = 0.012; *\*\*\*P* < 0.001. (**N**) The time until cells returned to baseline levels was determined. Mean ± SEM; neoAC CellC (n=3), AdAC CellC (n=4), AGES (n=4); ANOVA with Tukey’s post hoc test; P > 0.05.

### Transcriptomic profiling of astrocytes reveals major differences introduced by cell culturing, age and cell origin

To obtain a deeper understanding of the similarities and differences of directly isolated and cultured neonatal and adult astrocytes as well as AGES and NSCs, we performed an unbiased transcriptional profiling using RNA sequencing (RNA-seq). First, we aimed to delineate the differences we identified with regards to astrocyte marker expression as well as glial cell migration, calcium signaling and synapse pruning on the transcriptomic level. We therefore performed principal component analysis (PCA) on functionally grouped genes and confirmed key differences in astrocyte marker gene expression between cell types and culture conditions (Table S1, Fig. S3A), supplementing our analysis of astrocytic markers on the mRNA and protein levels (Fig. 1D-I). Additionally, we observed that gene signatures corresponding to glial cell migration (Fig. S3B) and calcium signaling (Fig. S3C) separated AGES from primary astrocytes, matching the observed functional differences in wound gap and calcium responses. PCA analysis using genes related to synapse pruning, revealed clustering of AGES and cultured astrocytes separately from NSCs, validating their similarities in the synapse uptake assay (Fig. S3D). This analysis of functionally annotated gene clusters also indicated differences between cultured and directly isolated astrocytes supporting the notion that the migratory and synapse pruning capacity of astrocytes change upon culturing.

To obtain an unbiased view upon the transcriptional differences between our various astrocyte populations, we performed a PCA using the complete transcriptomic signature of each cell type, which showed appreciable variation between cell types up to the fourth component (Fig. 3A-C). To functionally annotate the underlying genetic differences driving the PCA, we applied gene set enrichment analysis to the genes constituting the four PCs (Fig. 3D-E; Table S3 for complete gene set list). Additionally, differentially expressed genes between two sets of cell types were identified and functionally annotated using the R tmod package for gene set enrichment analysis ^44^ (Fig. 3F, Fig. S3F, Table S2 for complete gene set list).

**Figure 3:**
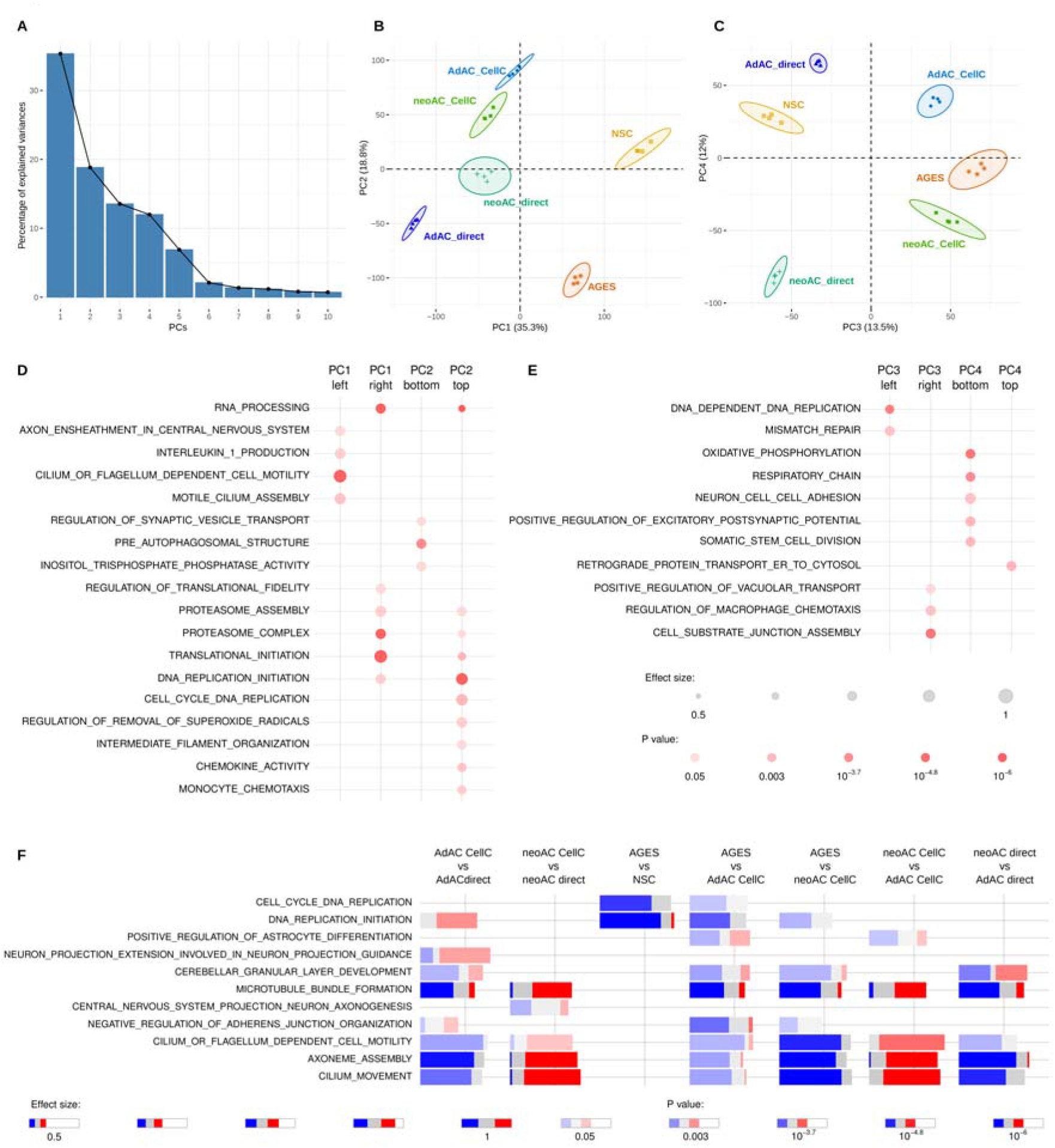
Transcriptomic profiling of astrocytes reveals major differences introduced by cell culturing, age and cell origin. RNA-seq was performed from cultured neonatal astrocytes (neoAC CellC; n=4), directly isolated neonatal astrocytes (neoAC direct; n=4), cultured adult astrocytes (AdAC CellC; n=4), directly isolated adult astrocytes (AdAC direct; n=4), AGES (n=4) and NSCs (n=4). (**A-C**) Principal component analysis (PCA) of RNA-seq results was performed and gene set enrichment analysis applied to genes ordered by their PCA loadings. The variance explained by the components is shown on the x- and y-axis. (**D-E**) Selected gene ontology (GO) terms characterizing each principal component are shown. The size of each dot reflects the effect size while the p-value is visualized by color intensity. (**F**) Differential gene expression-based functional enrichment analysis between cell types was done using the R tmod package. Each bar presents the fraction of significantly up (red) and downregulated (blue) genes in that particular GO category. The effect size is the area under the curve (see Fig. S3F).

Many studies neglect the comparison of stem-cell derived models to either neonatal or adult primary astrocytes and only show comparisons to ESCs and NSCs. The few studies that did compare iPSC-derived astrocytes to primary cells ^25,27^ did not perform global assessments of phenotypic, functional and transcriptional differences. Our transcriptomic analysis showed that primary astrocytes expressed genes involved in cilium and axoneme function (PC1 left), with NSCs and AGES being characterized by increases in translation initiation, RNA processing and proteasome activity (PC1 right) (Fig. 3B, D). tmod analysis comparing AGES to cultured primary cells identified the downregulation of genes related to DNA replication, cerebellar granular layer development and microtubule bundle formation and alterations in astrocyte cell differentiation genes (Fig. 3F). The negative regulation of adherens junction organization and cilium function (Fig. 3F) could explain the observed differences in closing wound gaps in AGES, as they are crucial for a rapid cell-cell contact remodeling during wounding ^28,29^. Underling our findings, PCA analysis using published transcriptomic data of other murine stem-cell derived astrocytes ^30^ revealed a clear separation from primary astrocytes indicating major gene expression changes between primary and stem cell-derived cells (Fig. S3E). The major gene expression and functional differences identified in our study indicate that AGES are not yet fully differentiated into astrocytes usually assisted and driven by the brain’s microenvironment and thus have not yet reached the functional and genetic profile of primary astrocytes. In order for stem-cell derived astrocytes to serve as a tool to study a broad range of astrocyte functions, we need to fully understand their similarities and differences to primary astrocytes and culture protocols need to be adapted to more closely match the profiles of primary cells.

Culturing of astrocytes induced upregulation of genes enriched in translation initiation, RNA processing, proteasome activity, chemotaxis and vesicle transport (PC2 top/PC3 right), and the downregulation of genes involved in autophagy, inositol lipid pathway and synaptic vesicle transport (PC2 bottom) and cell division (PC3 left, Fig. 3B-E). tmod analysis specifically revealed downregulation of cilium functions and microtubule formation and an upregulation of cell divison-related genes in adult astrocytes upon culturing. Neonatal astrocytes, on the other hand, upregulated microtubule formation and cilium functions *in vitro* (Fig. 3F). One of very few studies comparing directly isolated and cultured astrocytes found a downregulation of Wnt and Notch signaling upon culturing of neonatal astrocytes isolated by immunopanning ^11^. Interestingly, primary cilia alterations identified in our study are known to modulate the transduction of Wnt signaling ^31^. To our knowledge, alterations introduced by cell culturing of adult astrocytes were not assessed so far. In summary, the changes introduced *in vitro* thus reflect the change in microenvironment upon culturing in an age-dependent manner since the metabolic support of other cells is no longer required. AGES compared to NSCs downregulated genes indicative of cell cycle regulation and upregulated autophagy-related genes (PC2, Fig. 3B, D, F) which are common features of NSC differentiation ^32,33^ and indicates successful reprogramming into a new lineage with a modest proliferation behavior typical for astrocytes ^11^.

Age-specific differences included upregulated protein trafficking in adult astrocytes (PC4 top) and increased respiration, cell-cell adhesion, synapse regulation and stem cell division in neonatal astrocytes (PC4 bottom) (Fig. 3C, E). tmod analysis revealed that, *in vitro*, neonatal astrocytes show a pronounced upregulation of cilium functions and microtubule formation, compared to adult astrocytes. Directly isolated neonatal astrocytes showed alterations in genes linked to cerebellar granular layer development along with a downregulation in microtubule formation and cilium movement in contrast to directly isolated adult astrocytes (Fig. 3F). Thus, both the neonatal and astrocyte cultures retain key physiological functional characteristics carried out during postnatal neural development and aging respectively ^6^.

We identified striking differences in cilia function observed between primary astrocytes and AGES yet also between neonatal and adult astrocytes. Primary non-motile cilia have been described in both astrocytes ^34^ as well as human embryonic and neural stem cells ^35,36^ and are known to regulate cell division, neurogenesis, cell development and the response to certain extracellular signals ^34, 37, 38, 39^. Whilst the presence of primary cilia has not yet been described in stem-cell derived astrocytes, our results indicate that cilia properties in AGES compared to primary astrocytes are very different, yet also depend on maturation and culturing. These differential stages of cilia development and altered ciliogenesis in AGES, could also provide an explanation for their differential response to wound gaps. All in all, the transcriptomic differences we observed were driven by cell origin (primary vs. NSC-derived cells), culturing of cells and developmental stage of primary cells (Fig. S4).

In summary, our in-depth molecular side-by-side analysis of directly isolated and cultured mouse primary astrocytes and AGES showed clear differences in astrocyte marker profiles, functional readouts and transcriptomic signatures. Physiological *in vivo* functions driven by intercellular interactions are lost in cultured primary astrocytes. Although *in vitro* modelling of astrocytes is an inevitable tool which has been crucial for vital advances and discoveries, our data highlights the need to better mimic the CNS microenvironment *in vitro* when investigating trophic support provided by astrocytes. Generating a CNS microenvironment in culture might also assist in differentiating ESCs or iPSCs into mature astrocytes with closer similarity to primary astrocytes, since transcriptome analyses indicated that AGES are not yet fully differentiated. Our investigation is also aimed at supplementing future analyses and comparisons of astrocyte differentiation and culturing protocols also in the context of regional astrocytic differences. Further, our study highlights the importance of making informed decisions about the biological question under investigation and to match it to a suitable astrocyte isolation and cultivation protocol.

---

**Figure S1:**
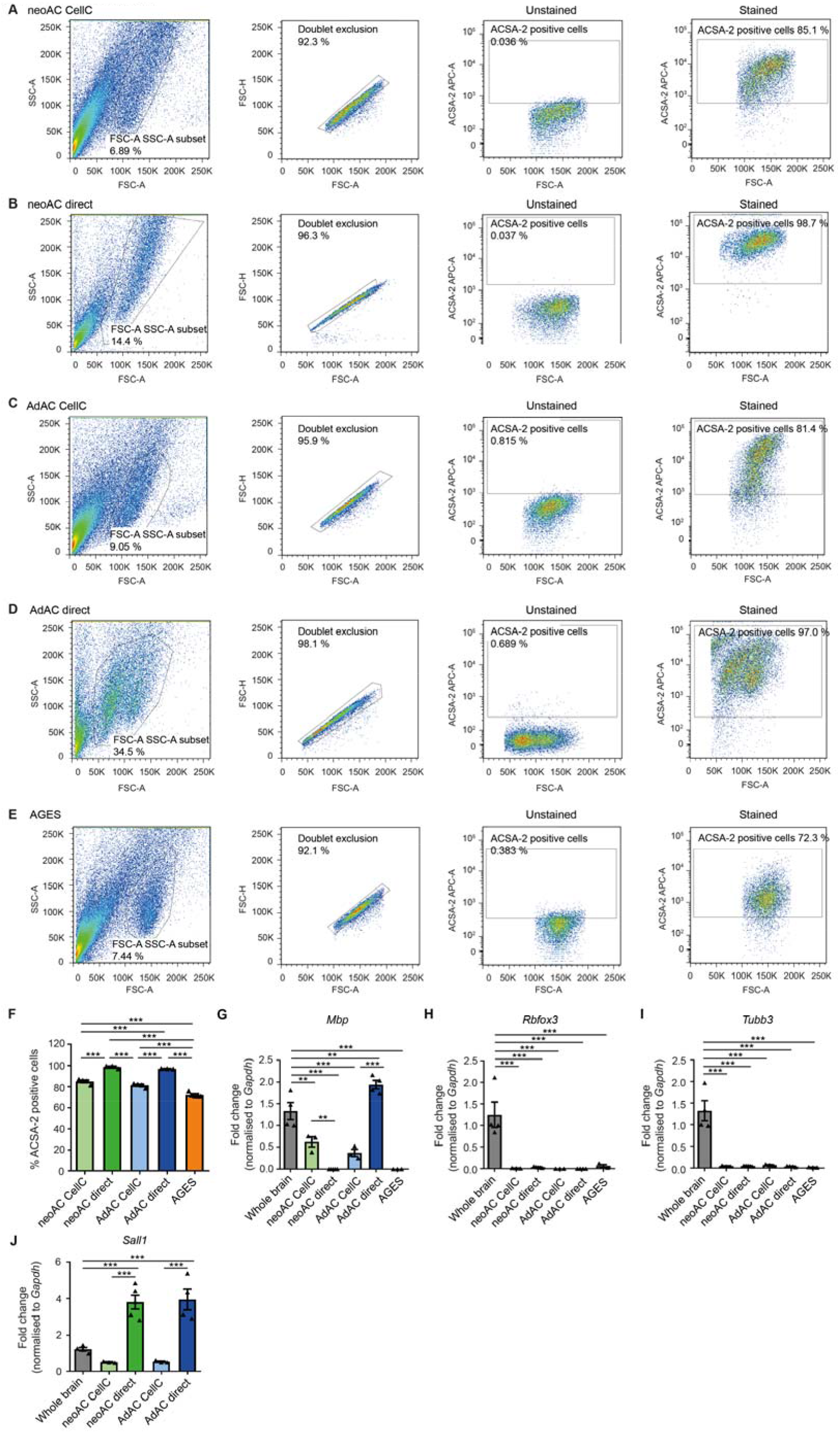
Cell purity and cell type composition changes during cell culturing. (**A-E**) Fluorescence activated cell sorting (FACS) of ACSA-2-labelled (stained) and unlabelled (unstained) cell suspensions is shown, demonstrating doublet exclusion and gating for APC-labelled ACSA-2 positive cells based on ACSA-2 negative stain in (**A**) cultured neonatal astrocytes (neoAC CellC), (**B**) directly isolated neonatal astrocytes (neoAC), (**C**) cultured adult astrocytes (AdAC CellC), (**D**) directly isolated adult astrocytes (AdAC) and (**E**) AGES. Representative FACS blots are shown for each cell type which include the average percentage of ACSA-2 positive cells. (**F**) Quantities of ACSA-2 expression levels in primary astrocytes and AGES as determined by FACS were summarized. Mean ± SEM; neoAC CellC (n=4), neoAC (n=3), AdAC CellC (n=4), AdAC (n=3), AGES (n=5). ANOVA with Tukey’s post hoc test *\*\*\*P* < 0.001 (**G-J**) Gene expression of non-astrocytic cell markers was determined by quantitative real-time PCR in cultured neonatal astrocytes (neoAC CellC), directly isolated neonatal astrocytes (neoAC), cultured adult astrocytes (AdAC CellC), directly isolated adult astrocytes (AdAC) and AGES. (**G**) *Mbp* was used as an oligodendrocytic marker (whole brain vs AdAC direct *\*\*P* = 0.008; whole brain vs neoAC CellC *\*\*P* = 0.004; *\*\*\*P* < 0.001). (**H**,**I**) *Rbfox3* (NeuN) and *Tubb3* represent neuronal marker genes (*\*\*\*P* < 0.001). (**J**) *Sall1* is a marker for microglia (*\*\*\*P* < 0.001). All expression values were normalized to the internal control *Gapdh* and whole brain lysates as a reference. Mean ± SEM; Whole brain (n=4), neoAC CellC (n=3), neoAC (n=5), AdAC CellC (n=3), AdAC (n=4), AGES (n=3); ANOVA with Tukey’s post hoc test; significances were only indicated for meaningful comparisons.

**Figure S2:**
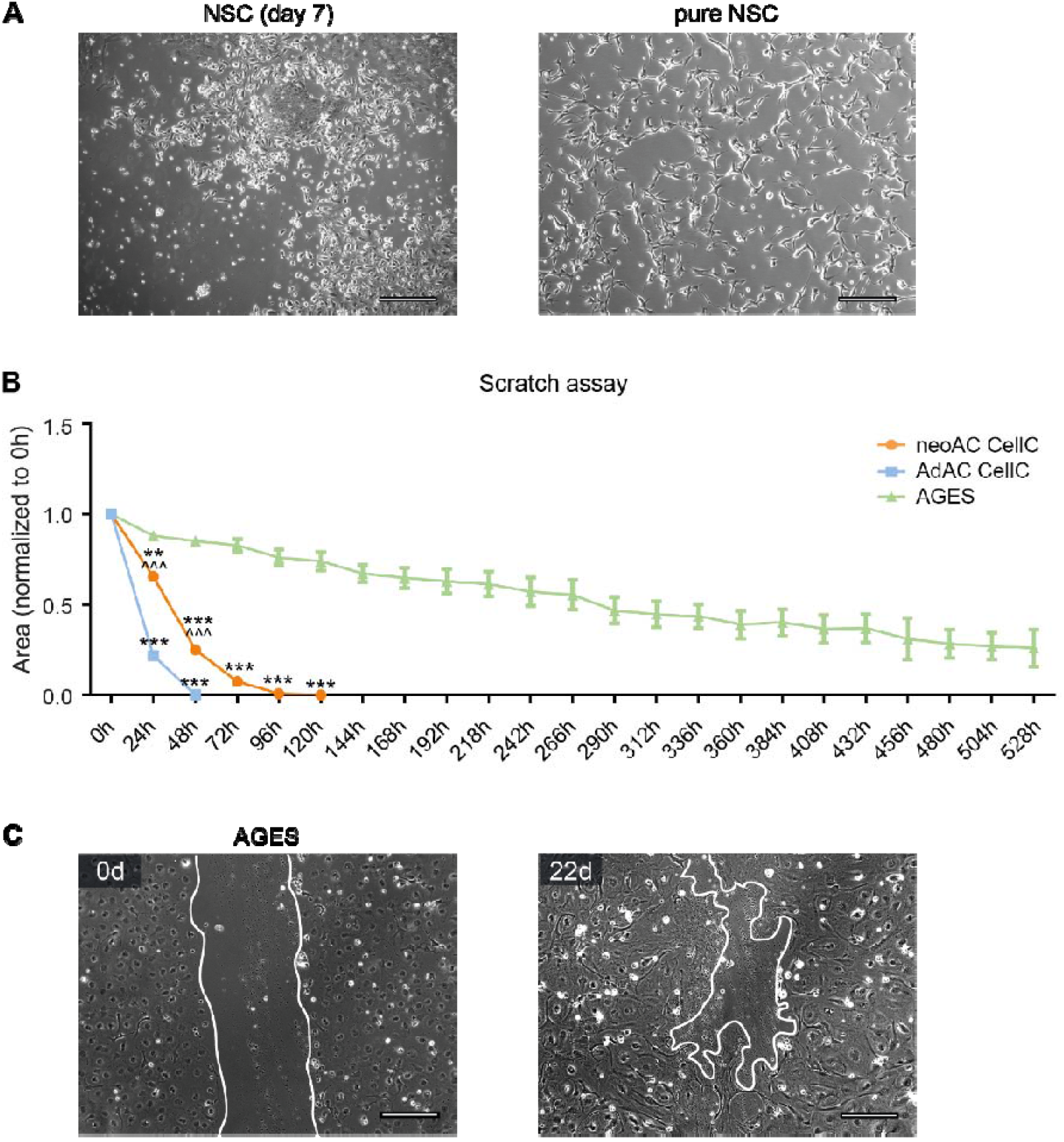
Migratory behavior of primary astrocytes and AGES over 22 days. (**A**) NSCs were differentiated from mESCs. For receiving pure bipolar NSC cultures, NSCs were cultured for at least eight passages in medium containing the growth factors EGF and FGF. Representative phase contrast images of NSC 7 days after starting the differentiation (mixed mESC and NSC culture) and pure NSC cultures are shown. Scale bar = 200 µm. (**B**) A scratch assay was used to determine the migratory behaviour of cultured neonatal astrocytes (neoAC CellC), cultured adult astrocytes (AdAC CellC) and AGES. The area of the wound gap normalized to time point 0 h is shown over time. * significance compared to AGES; ^ significance compared to AdAC CellC. Mean ± SEM; neoAC CellC (n=4), AdAC CellC (n=4), AGES (n=4); Two-way ANOVA with Bonferroni post hoc test; ***\*\*P*** = 0.004; *\*\*\*P* < 0.001; *^^^P* < 0.001. (**C**) Representative phase contrast images of the wound gap at the starting point (0 d) and after 22 days are shown of AGES. Scale bar = 200 µm.

**Figure S3:**
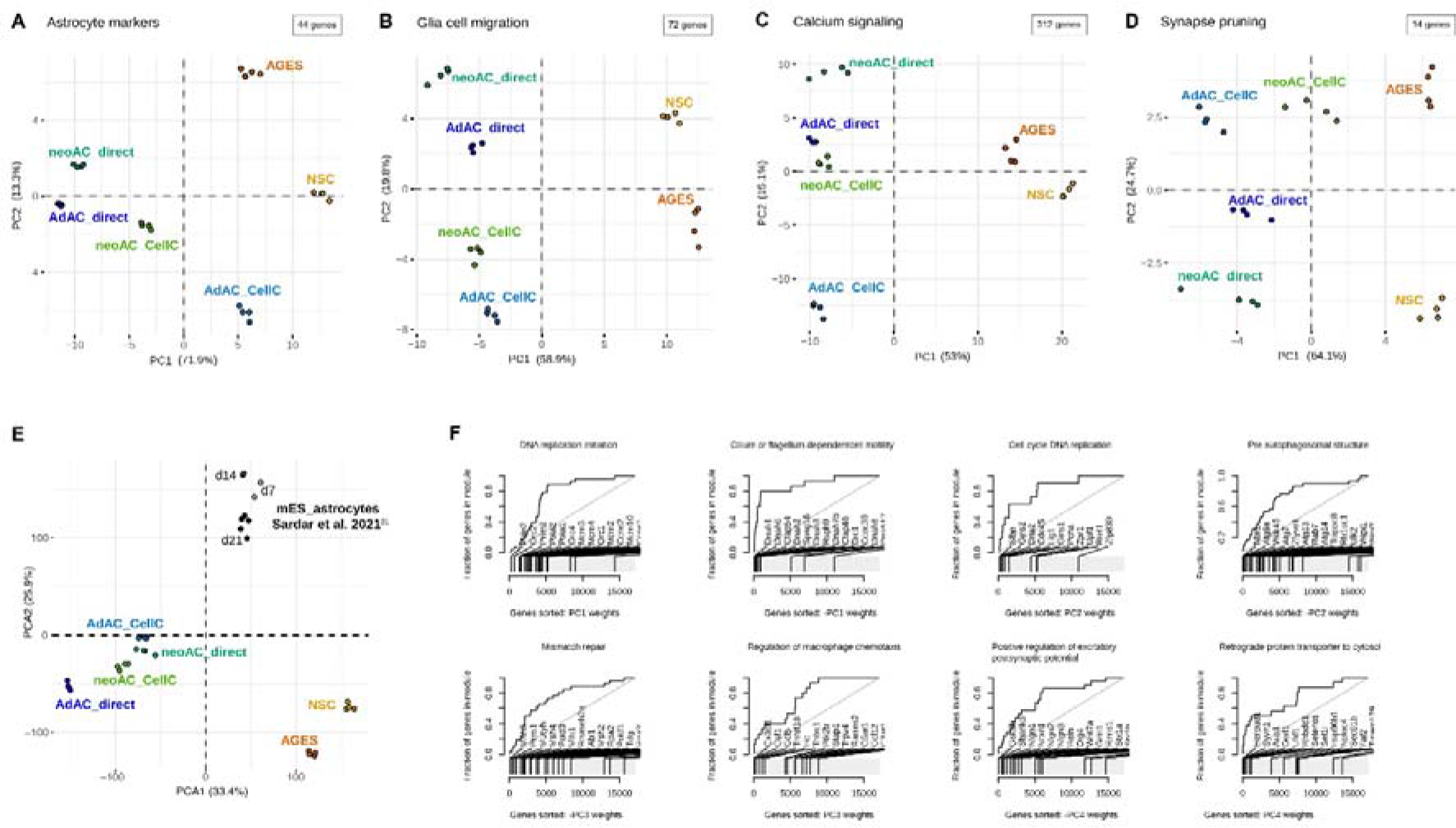
Principal component analysis and evidence plots of RNA-seq data underline functional differences between astrocytic cell types. RNA-seq was performed from cultured neonatal astrocytes (neoAC CellC; n=4), directly isolated neonatal astrocytes (neoAC direct; n=4), cultured adult astrocytes (AdAC CellC; n=4), directly isolated adult astrocytes (AdAC direct; n=4), AGES (n=4) and NSCs (n=4). (**A**) Principal component analysis (PCA) of published astrocytic marker genes ^5^ is shown. (**B-D**) Principal component analysis (PCA) of samples based only on genes from select GO terms relating to our functional assays was performed. The variance explained by the first and second component is shown on the x- and y-axis. (**E**) PCA of our samples and mouse embryonic stem cell-derived astrocytes (mES_astrocytes) from different time points of culture (day 7, 14, 21) published by Sardar et al. ^31^. (**F**) Evidence plots of select GO terms from Fig. 6B displaying genes sorted based on the gene PCA loadings on the x-axis and cumulative fraction of genes from a particular gene set (GO term) on the y-axis.

**Figure S4:**
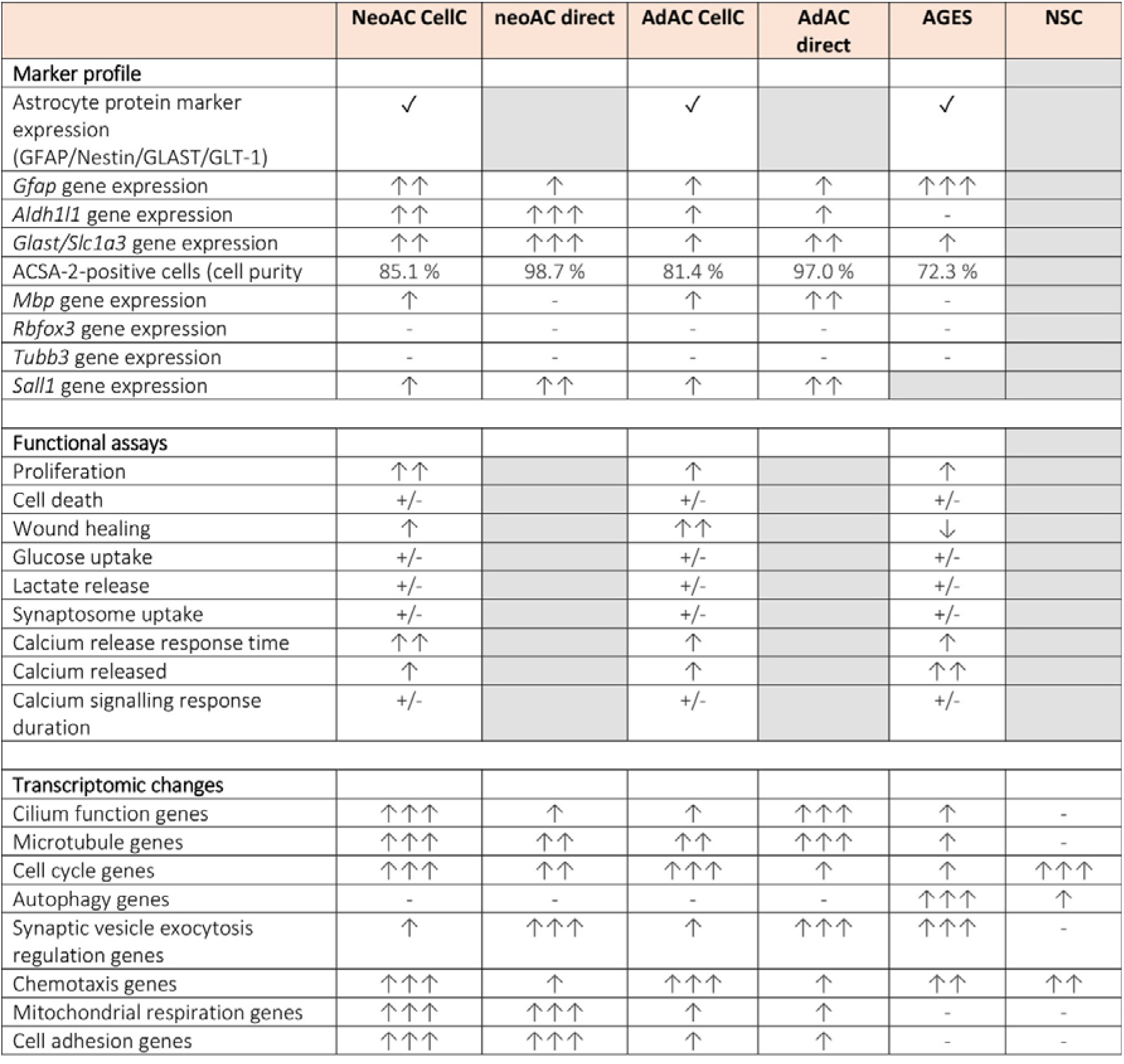
Summary of experimental results comparing cultured and directly isolated primary cells and AGES. Summary of the comparison of neonatal astrocytes (neoAC CellC), directly isolated neonatal astrocytes (neoAC direct), cultured adult astrocytes (AdAC CellC), directly isolated adult astrocytes (AdAC direct) and AGES. The following symbols were used for visualizing the obtained results: Grey box = The assay was not performed for these cells, ✓= presence for indicated attributes, ↑ = low expression/ functional activity, ↑↑ = intermediate expression/ functional activity, ↑↑↑ = high expression/ functional activity, - = no expression, +/- = no change in all analyzed groups.

## Methods

### Mice

For adult astrocyte cultures C57BL/6J mice at an age of > 100 days were used, whilst neonatal cells were taken from p 4-8 C57BL/6J pups. Female and male mice were mixed for all analyses. Mice were group housed under specific pathogen-free conditions on a 12 h light/dark cycle, and food and water were provided to the mice ad libidum. All animal experiments were performed in accordance with the national animal protection guidelines approved by the regional offices for health and social services in Berlin (LaGeSo, license numbers T 0276/07 and O298/17).

### Adult astrocyte isolation via MACS

Adult astrocytes were isolated using Magnetic-Activated Cell Sorting (MACS; Miltenyi Biotec) according to manufacturer’s protocol. In brief, mice were anesthetized with Isoflurane, euthanized with CO_2_, the brain dissected and placed in cold D-PBS supplemented with 0.91 mM CaCl_2_, 0.49 mM MgCl_2_-6H_2_O, 0.55 mM glucose and 0.033 mM sodium pyruvate (pH 7.2) (termed D-PBS). For direct astrocyte isolation, the olfactory bulb and cerebellum were removed and brains of two mice were pooled for tissue dissociation. Tissue dissociation was performed in C-tubes (Miltenyi, 130-096-334) using the Adult Brain Dissociation Kit (Miltenyi, 130-107-677) on program 37C_ABDK_01 of the gentleMACS™ Octo Dissociator with Heaters (Miltenyi Biotec, 130-096-427) allowing for simultaneous enzymatic and mechanical tissue disruption. Red blood cells and debris were removed and astrocytes magnetically labelled using ACSA-2 microbeads (Miltenyi Biotec, 130-097-678) according to manufacturer’s protocol. The resulting cell suspension was filtered once through a 70 µm pre-separation filter, consecutively passed through two MS columns (Miltenyi Biotec, 130-042-201) placed on a OctoMACS™ manual separator and flushed into Eppendorf tubes in 0.5 % BSA in PBS, pH 7.2 or cell culture medium (see section on astrocyte culture below). Cell pellets were collected, snap-frozen in liquid nitrogen and stored at −80 °C until further use.

### Neonatal astrocyte isolation via MACS

Neonatal astrocytes were isolated by using MACS from p 4-8 C57BL/6J mice. Cerebral tissue was isolated, meninges removed and brains of two mice pooled in a C-tube for tissue dissociation with the Neural Tissue Dissociation Kit (Miltenyi, 130-092-628) on program 37C_NTDK_1 of the gentleMACS™ Octo Dissociator with Heaters (Miltenyi Biotec, 130-096-427) allowing for simultaneous enzymatic and mechanical tissue disruption. Afterwards, the cell suspension was washed with Hanks’ Balanced Salt solution without calcium and magnesium (HBSS, Thermo Fisher, 14170138) and astrocytes magnetically labelled with ACSA-2 microbeads (Miltenyi Biotec, 130-097-678), according to manufacturer’s protocol. The resulting cell suspension was filtered once through a 70 µm pre-separation filter, consecutively passed through two MS columns (Miltenyi Biotec, 130-042-201) placed on a OctoMACS™ manual separator and flushed into Eppendorf tubes in 0.5 % BSA in PBS, pH 7.2 or cell culture medium (see section on astrocyte culture below). Cell pellets were collected, snap-frozen in liquid nitrogen and stored at −80 °C until further use.

### Adult and neonatal astrocyte culture

Cell culture plates (24-well or 96-well as indicated; BD Biosciences) were prepared two days prior to astrocyte isolation. First, wells were coated with 20 µg/ml poly-L-lysine (PLL) (Sigma, 2636-25MG) in PBS overnight at 37 °C, 5 % CO_2_. The next day, wells were washed twice with PBS and subsequently coated with 2 µg/ml laminin in PBS (Sigma, L2020) overnight at 37 °C, 5 % CO_2_. For immunofluorescence, cells were plated onto 13 mm coverslips (VWR) coated with 0.5 mg/ml PLL and 10 µg/ml laminin. For calcium imaging, cells were plated onto glass bottom dishes (MatTek, P35G-1.5-14-C) coated with 0.5 mg/ml PLL and 10 µg/ml laminin. Astrocyte isolation was performed as described above under sterile conditions. ACSA-2 labelled cells were flushed from the MS column with pre-warmed AstroMACS medium (Miltenyi Biotec, 130-117-031) supplemented 50 U/ml penicillin/streptomycin (Sigma, P0781-20ML) and 0.25 % L-glutamine (0.5 mM; Thermo Fisher, 25030-024). Neonatal and adult astrocytes were cultured using the same medium composition. Cells were plated at 100,000 cells/24 well or glass dish or 25,000 cells/96 well onto the middle of the well in a droplet of AstroMACS medium and incubated at 37 °C, 5 % CO_2_ for 1-3 h before filling up the medium. Medium was changed every three days and grown for 7-10 days before use.

### Astrocyte differentiation from embryonic stem cells

Astrocytes were differentiated from mouse embryonic stem cells (mESCs; GSC-5003, MTI-GlobalStem) using a previously published protocol^16^. mESCs were first differentiated into neural stem cells (NSCs) by plating mESCs as single cells on flasks coated with 0.1 % gelatin (Sigma, G9391-100G) in N2B27 medium (1 part DMEM/F12 medium (Thermo Fisher, 11320-033), 1 part Neurobasal medium (Thermo Fisher, 21103-049) supplemented with 1 % N2 supplement (Thermo Fisher, 17502-048), 2 % B27 supplement (Thermo Fisher, 17504-044), 2 mM Glutamax (Thermo Fisher, 35050-038), 55 µM 2-mercaptoethanol (Thermo Fisher, 21985-023), 0.075 % Insulin (Sigma, I0278-5ML), 50 mg/ml bovine serum albumin (BSA, Thermo Fisher, 15260-037) and 50 U/ml penicillin/streptomycin (Sigma, P0781-20ML). After differentiating mESCs to NSCs for 14 days, cells were detached with 0.05 % trypsin and re-plated on 0.1 % gelatin-coated flasks as single cells in N2B27 medium supplemented with 20 ng/ml FGF (Peprotech, 450-33) and 20 ng/ml EGF (Peprotech, AF-315-09). At this stage, the NSC culture consisted of an inhomogeneous culture with bipolar, triangular and aggregating cells (Fig. S2A). For selection and maintenance of bipolar NSCs, cells were split at a density of 80-90 % by tapping flasks for removing cell aggregates. Cells were washed with PBS once and detached with 0.05 % trypsin for 10-15 sec. Here, the more trypsin sensitive-bipolar NSCs went into suspension while triangular cells remained attached. After 10-15 sec of trypsinization, trypsin was diluted with 1 part PBS and the cell suspension was filtered through a 70 µm cell strainer into a tube prefilled with 2 parts PBS. Cells were centrifuged at 900 g for 3 min and resuspended in N2B27 medium supplemented with 20 ng/ml FGF and 20 ng/ml EGF. For all experiments described here, NSCs with a passage number from 50-60 were used to assure full purity.

For differentiating NSCs into astrocytes generated from ESCs (AGES), plates or dishes were coated with 10 µg/ml Poly-L-Ornithine (PLO, Sigma, P4957) in PBS for at least 2 h, wells were washed twice with PBS and subsequently coated with 2 µg/ml laminin (Sigma, L2020) in PBS overnight at 37 °C, 5 % CO_2_. NSCs were detached as described for cell maintenance and seeded at a density of 100,000 cells into precoated 24 wells in N2B27 medium supplemented with 20 ng/ml BMP4 (Peprotech, 315-27). After 3 days of differentiation in BMP4-supplemented N2B27 medium, AGES were used for described experiments. Medium was changed on day 1 of differentiation and every 3 days afterwards.

### Immunocytochemistry and confocal microscopy

Cells were washed once with pre-warmed PBS and fixed for 20 min at room temperature (RT) in freshly prepared 4 % paraformaldehyde (PFA) in PBS buffer, pH 7.4. Membranes were permeabilized with 0.1 % Triton-X100 in PBS for 20 min at RT and samples were blocked with freshly prepared 3 % bovine serum albumin (BSA, Merck) in PBS for 1 h at RT. GFAP (Abcam, ab4674, 1:1000), GLAST (Abcam, ab416, 1:200), Nestin (Abcam, Ab81755, 1:100) and GLT-1 (Sigma-Aldrich, Ab1783, 1:500) primary antibodies were diluted in PBS and incubated overnight at 4 °C. Following washes in PBS, Alexa Fluor^®^ 647 Goat Anti-Rabbit (Thermo Fisher, A21244), Alexa Fluor^®^ 647 Goat Anti-Guinea Pig (Thermo Fisher, A21450) and Cy™5 Donkey Anti-Chicken (Jackson ImmunoResearch, 703-175-155) secondary antibodies were diluted 1:500 in PBS and incubated at RT for 2 h. Nuclei were stained with 5 mg/ml 41,6-diamidino-2-phenylindole (DAPI; Roche, 1023627600) in PBS for 1 min and coverslips were mounted onto SuperFrost^®^ Plus slides (R. Langenbrink) with fluorescent Aqua-Poly/Mount mounting medium (Polysciences, 18606-20). The cells were imaged using a Leica TCS SP5 confocal laser scanning microscope and an HCX PL APO lambda blue 63× oil UV objective or HCX PL APO 40.0×1.30 OIL objective controlled by the LAS AF scan software (Leica Microsystem, Germany). Three-dimensional image stacks (1 µm step size) were taken and are shown as maximum projections.

### FACS analysis of purity

For Fluorescence-Activated Cell Sorting (FACS) of cultured cells, cells were detached in PBS using a cell scraper. For FACS analysis directly after cell isolation, cells were collected after flushing from MS columns. Cells were pelleted by centrifugation at 300 g, 10 min, 4 °C and stained with the APC-labelled ACSA-2 antibody (1:10, Miltenyi Biotec, 130-102-315) in 0.5 % BSA in PBS, pH 7.2 for 10 min at 4 °C. Cells were washed in 0.5 % BSA in PBS, pH 7.2 and collected by centrifugation followed by FACS analysis. Flow cytometry was performed using a FACSCanto II (BD Biosciences) and analyzed with FlowJo 7.6.5 software. Doublets were excluded by gating before analysing ACSA-2 levels.

### RNA isolation and quality control

Total RNA from frozen cell pellets or astrocytic cultures was isolated using the NucleoSpin miRNA kit (Macherey Nagel, 740971-250). RNA fractions containing small and large RNA were isolated and combined according to manufacturer’s instructions. For quantitative real-time PCR experiments, the rDNase treatment was omitted whilst for sequencing experiments, rDNase treatment was included. RNA was eluted using pre-warmed RNase-free H_2_O, snap-frozen in liquid nitrogen and stored at −80 °C until further use. RNA concentration was measured on a NanoQuant Plate™ using an Infinite^®^ 200 Pro plate reader and the i-control™ Microplate Reader Software (Tecan Life Sciences). For sequencing samples, RNA quality was further analysed by determining the RIN using the RNA ScreenTape (Aglient, 5067-5576) measured on the 4200 TapeStation system (Agilent). Samples with a RIN over 7 and a concentration of at least 100 ng total RNA were used.

### Quantitative real-time PCR

cDNA was synthesized from ≤ 1µg RNA using the QuantiTect Reverse Transcription Kit (Qiagen, 205311), snap-frozen and stored at −80 °C until further use. Gene expression analyses were performed on 12 ng cDNA per reaction using the TaqMan Fast Advanced Master Mix (ABI, 4364103) in a 384 well plate on a QuantStudio™ 6 Flex Real-Time PCR System (Thermo Fisher, A28139). Each PCR cycle had the following conditions for denaturation, annealing and lastly, extension: 95 °C for 20 sec 95 °C for 1 sec, and 60 °C for 20 sec. *Gapdh* was used as an internal control and the delta-delta Ct method was used for quantification. Three technical replicates per condition were performed. The following Taqman primers (Thermo Fisher) were used: *Gfap* (Mm01253033_m1), *Aldh1l1* (Mm03048957_m1), *Slc1a3* (Mm00600697_m1), *Mbp* (Mm01266402_m1), *Olig2* (Mm01210556_m1), *Sall1* (Mm00491266_m1), *Rbfox3* (Mm01248771_m1), beta-III-tubulin (*Tuj1*) (Mm00727586_s1), *Gapdh* (Mm99999915_g1).

### Quantitation of cell proliferation

Cell proliferation was measured based on the ability of living cells to incorporate EdU with the Click-iT™ EdU Microplate Assay (Invitrogen, 10214). The manufacturer’s protocol was followed. Briefly, cells cultured at a density of 50,000 cells per 96 well were incubated with 10 µM EdU diluted in respective medium for 72 h. Afterwards, the medium was removed and collected for the LDH cytotoxicity assay (see below) and cells were fixed and click-labeled. EdU was detected by using an anti-Oregon Green HRP antibody provided in the kit and incubated with an Amplex UltraRed reaction mixture for detection. Fluorescence was measured on the Infinite^®^ 200 Pro plate reader (Tecan Life Sciences) using an emission wavelength of 530 nm and an integration time of 40 µsec. The plate was read at the beginning of EdU incubation (t=0h) and after staining (t=72h) to normalize values for background fluorescence. Each measurement was normalized to reference absorbance at 600 nm. Two technical replicates per condition were performed.

### Cytotoxicity

Cytotoxicity was assessed by measuring lactate dehydrogenase (LDH), an enzyme released upon cell lysis, using the CytoTox 96^®^ Non-Radioactive Cytotoxicity Assay (Promega, G1780). As positive control for 100 % cytotoxicity, cells were lysed by adding 10 % Triton-X to the medium and incubated for 30 min at 37 °C. For LDH detection, the manufacturer’s protocol was followed. In brief, cell medium of astrocytes used for the cell proliferation assay (see above) was added to the CytoTox 96^®^ Reagent for 30 min before measuring absorbance at 492 nm (600 nm reference value) on an Infinite^®^ 200 Pro plate reader. All values were presented relative to lysed cell positive control. Two technical replicates per condition were performed.

### Wound sensing assay

A “wound gap” was created by scratching a straight line through the confluent monolayer of 100,000 cells/well in a 24 well plate using a sterile 200 µl pipette tip. To guarantee imaging of the same spot, a line was drawn on the outside of the well perpendicular to the scratch. The intersection was used as the area to take all photos. Photos were taken with a Carl Zeiss Axio observer Z1 inverted microscope at time point 0 h and every consecutive 24 h to monitor cellular migration. Quantification was performed using the ImageJ (NIH, Maryland, USA) ROI tool and the area which was not covered by cells was measured and normalized to the non-treated control.

### Glucose Uptake

Glucose uptake in AGES, neonatal and adult astrocytes plated on 96-well plates (50,000 cells per well) was measured with the Glucose Uptake-Glo™ Assay (Promega, J1341) according to manufacturer’s protocol. In brief, medium was removed and collected for the Lactate-Glo™ Assay (see below). Cells were washed once with PBS and 1 mM 2-deoxyglucose (2DG) in PBS was added for 10 min to induce glucose uptake. The reaction was stopped using Stop Buffer followed by the addition of Neutralisation Buffer and the 2DG6P Detection Reagent. After 2 h, luminescence was measured on the Infinite^®^ 200 Pro plate reader (Tecan Life Sciences) using no attenuation, 1000 msec integration time and 0 msec settle time. Each measurement was normalized to reference absorbance at 600 nm at 0 h. Wells not containing any cells but included in the measurement protocol were used as a background reference. A standard curve ranging from 2.5 µM to 20 µM was generated with the provided 2DG6P standard to determine the glucose uptake concentration per well. Two technical replicates per condition were performed.

### Lactate Release

Lactate release was measured on cell medium collected from AGES, neonatal and adult astrocytes using the Lactate-Glo™ Assay (Promega, J5021). An equal volume of Lactate Detection Reagent was added and the plate shaken for 60 sec. After 1 h, luminescence was measured on the Infinite^®^ 200 Pro plate reader (Tecan Life Sciences) using no attenuation, 1000 msec integration time and 0 msec settle time. Each measurement was normalized to reference absorbance at 600 nm and to cell-free medium as a background reference. A standard curve ranging from 3.125 µm to 200 µM was generated with the provided lactate standard and used to determine the lactate release concentration. Two technical replicates per condition were performed.

### Synaptosome uptake assay

For the analysis of synaptosome uptake by primary astrocytes and AGES, pH-sensitive labelled synaptosomes were used. Synaptosomes were isolated as previously described ^40^. Unperfused brains of C57BL/6J mice at 65-75 days were placed in half-frozen PBS and cortices removed. Cortices including hippocampus were cut into small pieces and homogenized with a Teflon Homogenizer in homogenization buffer (10.9 % sucrose, 20 mM HEPES, 0.029 % EDTA, protease inhibitor, pH 7.4). Afterwards, the homogenized brain was centrifuged for 10 min at 3000 rpm at 4 °C. The synaptosomes in the supernatant were pelleted for another 15 min at 12500 g at 4 °C. The pellet containing the synaptosomes was homogenized in homogenization buffer using the Teflon homogenizer. Afterwards, synaptosomes were separated using a gradient of 0.8 M sucrose layered by 1.2 M sucrose and ultracentrifugation at 20100 g for 1 h 10 min at 4 °C. The synaptosome band formed between both sucrose layers was collected. Synaptosomes were pelleted by centrifugation at 12500 g for 15 min and resuspended in 0.1 M sodium bicarbonate to label them with a pH-sensitive fluorogenic dye (pHrodo™ Red succinimidyl ester, Thermo Fisher, 10676983, 1:660 dilution) by rotation for 2 h at RT. After washing labelled synaptosomes three times, they were resuspended in PBS containing 5 % DMSO. Synaptosomes from one brain were resuspended in 100 µl final volume and were diluted 1:12.5 in culture medium and added to the cells for 24 h before reading fluorescence with an Infinite^®^ 200 Pro plate reader at 560 nm excitation wavelength/ 585 nm emission wavelength.

### Calcium imaging

Calcium imaging was performed on cells grown on glass bottom dishes (35 mm dish, 14 mm glass; MatTek, 35G-1.5-14-C) using the Fluo-4, AM calcium indicator (Thermo Fisher, F14201). For reconstitution, a 20 % Pluronic^®^ F-127 (Sigma, P2443) in DMSO solution was mixed 1:1 with 10 mM Fluo-4AM in DMSO to yield a 10 % Pluronic^®^ F-127/5 mM Fluo-4, AM working solution. On the day of imaging, cells were incubated with 5 µM Fluo-4, AM solution dissolved in sterile HEPES buffer (150 mM NaCl, 5.4 mM KCl, 1.3 mM CaCl_2_, 0.83 mM MgSO_4_ x 7 H_2_O, 10 mM HEPES, 5 mM D(+)-Glucose, pH 7.4) for 30 min at RT. Afterwards, the Fluo4, AM was removed and cells kept in HEPES buffer at 37 °C, 5 % CO_2_ until and during imaging. The dish was transferred to an Okolab Incubation Chamber. Imaging was conducted at the Advanced Medical Bioimaging Core Facility (AMBIO) at the Charité Universitätsmedizin Berlin using the Nikon Spinning Disc Confocal CSU-X setup with the Nikon Eclipse TI microscope. An ATP solution for inducing calcium release in astrocytes was prepared with a final concentration of 100 µM in HEPES buffer. Using the peristaltic perfusion system (Multi Channel Systems, PPS5), the cells were perfused with HEPES for 2 min, followed by a 30 sec perfusion with 100 µM ATP and finally 1 min 30 sec in HEPES with a flow rate of 2.5 ml/min. Imaging was started after perfusing cells for 1 min 50 sec with HEPES. Intracellular Ca^2+^ changes were detected using the NIS Elements Imaging Software (Nikon, Version 5.10). Analysis was conducted using the ImageJ software with the Time Series Analyzer V3 plugin. The magnitude of Ca^2+^ concentration changes was detected via a temporal analysis of 10 single cells per n as a fluorescence intensity ratio F/F0. F0 was determined at a baseline 10 second window prior to ATP administration. For analysis, the average maximum fluorescence intensity, the time until cells responded to ATP treatment and the peak width (time between maximum intensity and reaching basal levels) was determined.

### Library preparation and RNA sequencing

RNA sequencing libraries were prepared using the TruSeq™ Stranded mRNA Library Prep kit (20020594, Illumina) starting from 100 ng of total RNA (RIN ≥7) on an ep*Motion*^®^ 5075 TMX workstation (Eppendorf). Library QC included size distribution check (BioAnalyser) and concentration determination with KAPA Library Quantification Kit (KK4857, Roche). Libraries were equimolar pooled and loaded on Illumina NovaSeq 6000 SP flowcell at 300 pM loading concentration with 1 % of PhiX by-mix. Sequencing was performed in paired-end 2×100 nt sequencing mode with 8nt index (i7) read.

### Data analysis

RNA-seq reads were mapped to mouse genome (GRCm38/p5) with STAR (version 2.7.3a) using the following parameters: --outFilterType BySJout --outFilterMultimapNmax 20 --alignSJoverhangMin 8 -

-alignSJDBoverhangMin 1 --outFilterMismatchNmax 999 --outFilterMismatchNoverLmax 0.04 -- alignIntronMin 20 --alignIntronMax 1000000 --alignMatesGapMax 1000000 ^41^. We obtained on average 87.7% uniquely mapped reads per sample. Reads were assigned to genes with *featureCounts* (-t exon -g gene_id, version 2.0.1) using Gencode GRCm38/vM12 gene annotation ^42^. The differential expression analysis was carried out with DESeq2 (version 1.22.1) using default parameters. For the principal component analysis (PCA) (in Fig.6 B) we used rlog (DESeq2, version1.22.1) transformed counts ^43^. Gene ontology enrichment analysis was done with CERNO algorithm from R tmod (0.46.2) package ^44^: for two sided comparisons (eg. AdAC CellC vs direct) genes were sorted by their adjusted (Benjamini Hochberg) p-values, and for PCA-based enrichment analysis genes were sorted by their PCA loadings. The enrichment analysis was done using GO gene set collection from MsigDB (r msigdbr 6.2.1). Subcategories BP, MF and CC were combined for the analysis.

### Statistics

Data were generated as multiple exploratory analyses to generate hypotheses and biostatistical planning for future confirmatory studies. Data analysis, processing, descriptive and formal statistical testing were done according to the current customary practice of data handling using Excel 2016, GraphPad PRISM 5.0 and ImageJ. All data generated or analysed during this study are included in this article.

Values are presented as mean ± SEM (standard error of the mean). Statistical difference between means was assessed either by the two-tailed t-test for two groups or ANOVA with the indicated post hoc test for more than two groups using the GraphPad Prism software. Outliers were not excluded and statistically significant values are indicated as *\*P* ≤ 0.05, *\*\*P* ≤ 0.01, and *\*\*\*P* ≤ 0.001.

## Acknowledgements

This project has received funding from the Deutsche Forschungsgemeinschaft (DFG, German Research Foundation) under Germany’s Excellence Strategy – EXC-2049 – 390688087, as well as HE 3130/6-1 to F.L.H., the German Center for Neurodegenerative Diseases (DZNE) Berlin, and the Innovative Medicines Initiative 2 Joint Undertaking under grant agreement No 115976. This Joint Undertaking receives support from the European Union’s Horizon 2020 research and innovation programme and EFPIA. S.S. was funded by a PhD fellowship of the NeuroCure Excellence Cluster EXC-2049.

## Author Contributions

KF and PE performed experiments and analysed data; AI and DB performed and supervised bioinformatical analyses; SS isolated synaptosomes; TB and S. Sauer performed and supervised RNA library preparation and RNA sequencing; FLH designed and supervised the study; KF, PE and AI prepared figures. All authors wrote, revised and approved the manuscript.

## Declaration of Interests

The authors declare that they have no conflict of interest.

## Notes

### Competing Interest Statement

The authors have declared no competing interest.

## References

1. Binmöller, F.-J. & Müller, C. M. (1992). Postnatal development of dye-coupling among astrocytes in rat visual cortex. Glia 6, 127–137.

2. Finkbeiner, S. (1992). Calcium waves in astrocytes-filling in the gaps. Neuron 8, 1101–1108.

3. Gordon, G. R., Choi, H. B., Rungta, R. L., Ellis-Davies, G. C. & Macvicar, B. A. (2008). Brain metabolism dictates the polarity of astrocyte control over arterioles. Nature 456, 745–749.

4. Rouach, N., Koulakoff, A., Abudara, V., Willecke, K. & Giaume, C. (2008). Astroglial metabolic networks sustain hippocampal synaptic transmission. Science 322, 1551–1555.

5. Araque, A., Carmignoto, G., Haydon, P. G., Oliet, S. H., Robitaille, R. & Volterra, A. (2014). Gliotransmitters travel in time and space. Neuron 81, 728–739.

6. Palmer, A. L. & Ousman, S. S. (2018). Astrocytes and Aging. Front Aging Neurosci 10, 337.

7. Matias, I., Morgado, J. & Gomes, F. C. A. (2019). Astrocyte Heterogeneity: Impact to Brain Aging and Disease. Front Aging Neurosci 11, 59.

8. Guttenplan, K. A. & Liddelow, S. A. (2019). Astrocytes and microglia: Models and tools. J Exp Med 216, 71–83.

9. McCarthy, K. D. & De Vellis, J. (1980). Preparation of separate astroglial and oligodendroglial cell cultures from rat cerebral tissue. J Cell Biol 85, 890–902.

10. Zhang, Y., Sloan, S. A., Clarke, L. E., Caneda, C., Plaza, C. A., Blumenthal, P. D., Vogel, H., Steinberg, G. K., Edwards, M. S., Li, G., Duncan, J. A., 3rd, Cheshier, S.H., Shuer, L. M., Chang, E. F., Grant, G. A., Gephart, M. G. & Barres, B. A. (2016). Purification and Characterization of Progenitor and Mature Human Astrocytes Reveals Transcriptional and Functional Differences with Mouse. Neuron 89, 37–53.

11. Foo, L. C., Allen, N. J., Bushong, E. A., Ventura, P. B., Chung, W. S., Zhou, L., Cahoy, J. D., Daneman, R., Zong, H., Ellisman, M. H. & Barres, B. A. (2011). Development of a method for the purification and culture of rodent astrocytes. Neuron 71, 799–811.

12. Rusnakova, V., Honsa, P., Dzamba, D., Ståhlberg, A., Kubista, M. & Anderova, M. (2013). Heterogeneity of Astrocytes: From Development to Injury – Single Cell Gene Expression. PLOS ONE 8, e69734.

13. Cahoy, J. D., Emery, B., Kaushal, A., Foo, L. C., Zamanian, J. L., Christopherson, K. S., Xing, Y., Lubischer, J. L., Krieg, P. A., Krupenko, S. A., Thompson, W. J. & Barres, B. A. (2008). A transcriptome database for astrocytes, neurons, and oligodendrocytes: a new resource for understanding brain development and function. J Neurosci 28, 264–278.

14. Batiuk, M. Y., De Vin, F., Duque, S. I., Li, C., Saito, T., Saido, T., Fiers, M., Belgard, T. G. & Holt, M. G. (2017). An immunoaffinity-based method for isolating ultrapure adult astrocytes based on ATP1B2 targeting by the ACSA-2 antibody. J Biol Chem 292, 8874–8891.

15. Kantzer, C. G., Boutin, C., Herzig, I. D., Wittwer, C., Reiss, S., Tiveron, M. C., Drewes, J., Rockel, T. D., Ohlig, S., Ninkovic, J., Cremer, H., Pennartz, S., Jungblut, M. & Bosio, A. (2017). Anti-ACSA-2 defines a novel monoclonal antibody for prospective isolation of living neonatal and adult astrocytes. Glia 65, 990–1004.

16. Kleiderman, S., Sa, J. V., Teixeira, A. P., Brito, C., Gutbier, S., Evje, L. G., Hadera, M. G., Glaab, E., Henry, M., Sachinidis, A., Alves, P. M., Sonnewald, U. & Leist, M. (2016). Functional and phenotypic differences of pure populations of stem cell-derived astrocytes and neuronal precursor cells. Glia 64, 695–715.

17. Lattke, M., Goldstone, R., Ellis, J. K., Boeing, S., Jurado-Arjona, J., Marichal, N., MacRae, J. I., Berninger, B. & Guillemot, F. (2021). Extensive transcriptional and chromatin changes underlie astrocyte maturation in vivo and in culture. Nat Commun 12, 4335.

18. Eng, L. F., Ghirnikar, R. S. & Lee, Y. L. (2000). Glial fibrillary acidic protein: GFAP-thirty-one years (1969-2000). Neurochem Res 25, 1439–1451.

19. Escartin, C., Galea, E., Lakatos, A. et al. (2021). Reactive astrocyte nomenclature, definitions, and future directions. Nat Neurosci 24, 312–325.

20. Roybon, L., Lamas, N. J., Garcia, A. D., Yang, E. J., Sattler, R., Lewis, V. J., Kim, Y. A., Kachel, C. A., Rothstein, J. D., Przedborski, S., Wichterle, H. & Henderson, C. E. (2013). Human stem cell-derived spinal cord astrocytes with defined mature or reactive phenotypes. Cell Rep 4, 1035–1048.

21. Guizzetti, M., Kavanagh, T. J. & Costa, L. G. (2011). Measurements of astrocyte proliferation. Methods Mol Biol 758, 349–359.

22. Deitmer, J. W., Theparambil, S. M., Ruminot, I., Noor, S. I. & Becker, H. M. (2019). Energy Dynamics in the Brain: Contributions of Astrocytes to Metabolism and pH Homeostasis. Front Neurosci 13, 1301.

23. Chung, W. S., Clarke, L. E., Wang, G. X., Stafford, B. K., Sher, A., Chakraborty, C., Joung, J., Foo, L. C., Thompson, A., Chen, C., Smith, S. J. & Barres, B. A. (2013). Astrocytes mediate synapse elimination through MEGF10 and MERTK pathways. Nature 504, 394–400.

24. Shigetomi, E., Kracun, S., Sofroniew, M. V. & Khakh, B. S. (2010). A genetically targeted optical sensor to monitor calcium signals in astrocyte processes. Nat Neurosci 13, 759–766.

25. Julia, T.W.C., Wang, M., Pimenova, A. A., Bowles, K. R., Hartley, B. J., Lacin, E., Machlovi, S. I., Abdelaal, R., Karch, C. M., Phatnani, H., Slesinger, P. A., Zhang, B., Goate, A. M. & Brennand, K. J. (2017). An Efficient Platform for Astrocyte Differentiation from Human Induced Pluripotent Stem Cells. Stem Cell Rep 9, 600–614.

26. Bardehle, S., Krüger, M., Buggenthin, F., Schwausch, J., Ninkovic, J., Clevers, H., Snippert, H. J., Theis, F. J., Meyer-Luehmann, M., Bechmann, I., Dimou, L & Götz, M. (2013). Live imaging of astrocyte responses to acute injury reveals selective juxtavascular proliferation. Nat Neurosci 16, 580–586.

27. Bradley, R. A., Shireman, J., Mcfalls, C., Choi, J., Canfield, S. G., Dong, Y., Liu, K., Lisota, B., Jones, J. R., Petersen, A., Bhattacharyya, A., Palecek, S. P., Shusta, E. V., Kendziorski, C. & Zhang, S.-C. (2019). Regionally specified human pluripotent stem cell-derived astrocytes exhibit different molecular signatures and functional properties. Development 146, dev170910.

28. Collinet, C. & Lecuit, T. (2013). Stability and dynamics of cell-cell junctions. Prog Mol Biol Transl Sci 116, 25–47.

29. Veland, I. R., Lindbæk, L. & Christensen, S. T. (2014). Linking the Primary Cilium to Cell Migration in Tissue Repair and Brain Development. BioScience 64, 1115–1125.

30. Sardar D., Lozzi B., Woo J., Huang T. W., Cvetkovic C., Creighton C. J., Krencik R. & Deneen B. (2021) Mapping Astrocyte Transcriptional Signatures in Response to Neuroactive Compounds. Int J Mol Sci 22, 3975.

31. Gerdes, J.M. & Katsanis, N (2008). Ciliary function and Wnt signal modulation. Curr Top Dev Biol 85, 175–95.

32. Zhang, J., Nuebel, E., Daley, G. Q., Koehler, C. M. & Teitell, M. A. (2012). Metabolic regulation in pluripotent stem cells during reprogramming and self-renewal. Cell Stem Cell 11, 589–595.

33. Pan, H., Cai, N., Li, M., Liu, G.-H. & Izpisua Belmonte, J. C. (2013). Autophagic control of cell ‘stemness’. EMBO Mol Med 5, 327–331.

34. Sterpka, A., Yang, J., Strobel, M., Zhou, Y., Pauplis, C. & Chen, X. (2020). Diverged morphology changes of astrocytic and neuronal primary cilia under reactive insults. Mol Brain 13, 28.

35. Kiprilov, E. N., Awan, A., Desprat, R., Velho, M., Clement, C. A., Byskov, A. G., Andersen, C. Y., Satir, P., Bouhassira, E. E., Christensen, S. T., & Hirsch, R. E. (2008). Human embryonic stem cells in culture possess primary cilia with hedgehog signaling machinery. J Cell Biol 180, 897–904. https://doi.org/10.1083/jcb.200706028

36. Tong, C. K., Han, Y. G., Shah, J. K., Obernier, K., Guinto, C. D., & Alvarez-Buylla, A. (2014). Primary cilia are required in a unique subpopulation of neural progenitors. PNAS 111, 12438–12443.

37. Breunig, J. J., Sarkisian, M. R., Arellano, J. I., Morozov, Y. M., Ayoub, A. E., Sojitra, S., Wang, B., Flavell, R. A., Rakic, P. & Town, T. (2008). Primary cilia regulate hippocampal neurogenesis by mediating sonic hedgehog signaling. PNAS 105, 13126–13131.

38. Rohatgi, R., Milenkovic, L. & Scott, M. P. (2007). Patched1 regulates hedgehog signaling at the primary cilium. Science 317, 372–376.

39. Yoshimura, K., Kawate, T. & Takeda, S. (2011). Signaling through the primary cilium affects glial cell survival under a stressed environment. Glia 59, 333–344.

40. Liu, C., Kershberg, L., Wang, J., Schneeberger, S. & Kaeser, P. S. (2018). Dopamine Secretion Is Mediated by Sparse Active Zone-like Release Sites. Cell 172, 706-718.e715.

41. Dobin, A., Davis, C. A., Schlesinger, F., Drenkow, J., Zaleski, C., Jha, S., Batut, P., Chaisson, M. & Gingeras, T. R. (2012). STAR: ultrafast universal RNA-seq aligner. Bioinformatics 29, 15–21.

42. Liao, Y., Smyth, G. K. & Shi, W. (2014). featureCounts: an efficient general purpose program for assigning sequence reads to genomic features. Bioinformatics 30, 923–930.

43. Love, M. I., Huber, W. & Anders, S. (2014). Moderated estimation of fold change and dispersion for RNA-seq data with DESeq2. Genome Biology 15, 550.

44. Zyla, J., Marczyk, M., Domaszewska, T., Kaufmann, S. H. E., Polanska, J. & Weiner, J., 3rd (2019). Gene set enrichment for reproducible science: comparison of CERNO and eight other algorithms. Bioinformatics 35, 5146–5154.

